# Arabidopsis CALMODULIN-BINDING PROTEIN 60b plays dual roles in plant immunity

**DOI:** 10.1101/2021.04.15.440066

**Authors:** Weijie Huang, Zhongshou Wu, Hainan Tian, Xin Li, Yuelin Zhang

## Abstract

Arabidopsis SYSTEMIC ACQUIRED RESISTANCE DEFICIENT 1 (SARD1) and CALMODULIN-BINDING PROTEIN 60g (CBP60g) are two master transcription factors that regulate many defense-related genes in plant immunity. They are required for immunity downstream of the receptor-like protein SUPPRESSOR OF NPR1-1, CONSTITUTIVE 2 (SNC2). Constitutive defense responses in the gain-of-function autoimmune *snc2-1D* mutant are modestly affected by either *sard1* or *cbp60g* single mutants, but completely suppressed by the *sard1 cbp60g* double mutant. Here we report that CBP60b, another member of the CBP60 family, also functions as a positive regulator of SNC2-mediated immunity. Loss-of-function mutations of *CBP60b* suppress the constitutive expression of *SARD1* and enhanced disease resistance in *cbp60g-1 snc2-1D*, whereas over-expression of *CBP60b* leads to elevated *SARD1* expression and constitutive defense responses. In addition, transient expression of *CBP60b* in *Nicotiana benthamiana* activates the expression of the *pSARD1::luciferase* reporter gene. Chromatin immunoprecipitation assay further showed that CBP60b is recruited to the promoter region of *SARD1*, suggesting that it directly regulates *SARD1* expression. Interestingly, knocking out *CBP60b* in the wild type background leads to ENHANCED DISEASE SUSCEPTIBILITY 1 (*EDS1*)-dependent autoimmunity, suggesting that CBP60b is required for the expression of a guardee/decoy or a negative regulator in immunity mediated by receptors carrying an N-terminal TIR (Toll-interleukin-1 receptor-like) domain.

**Significance statement:** Arabidopsis SARD1 serves as a master transcription factor in plant immunity. In this study, we showed that CBP60b positively regulates *SARD1* expression, and TIR signaling is activated when CBP60b is inactivated.

## Introduction

Plants have evolved sophisticated innate immune systems to fight against various pathogens (Jones and Dangl 2006, Zhou and Zhang, 2020). Upon infection, plasma membrane-localized pattern recognition receptors (PRRs) recognize pathogen/microbe-associated molecular patterns (PAMPs or MAMPs), which are conserved molecules associated with groups of pathogens, to initiate pattern-triggered immunity (PTI)(Monaghan and Zipfel, 2012). To facilitate colonization, pathogens have evolved various effector proteins to interfere with PTI (Dou and Zhou, 2012). Meanwhile, plants evolved resistance (R) proteins, mostly in the nucleotide-binding leucine-rich repeat (NLR) protein family, to perceive the pathogen effectors and activate effector-triggered immunity (ETI)(van Wersch *et al*., 2020). Activation of PTI and ETI at local infection sites further leads to development of systemic acquired resistance (SAR) in the distal parts of the plants (Sun and Zhang, 2021).

One class of plant PRRs belong to the receptor-like protein (RLP) family. RLPs usually have a short cytoplasmic tail, a single transmembrane motif and an extracellular leucine-rich repeats (LRR) domain involved in binding to ligands. One of the *Arabidopsis* RLPs, SNC2 (SUPPRESSOR OF NPR1-1, CONSTITUTIVE 2), contributes to defense against bacteria (Zhang *et al*., 2010b). On one hand, loss of SNC2 results in enhanced susceptibility to bacterial pathogen *Pseudomonas syringae* pv *tomato* (*Pto*) DC3000. On the other hand, a gain-of-function mutation in *snc2-1D* results in constitutive *Pathogenesis-Related* (*PR*) gene expression, elevated levels of the plant defense hormone SA, and enhanced resistance against the oomycete pathogen *Hyaloperonospora arabidopsidis* (*Hpa*) Noco2 (Zhang *et al*., 2010b). Due to constitutively activated immunity, *snc2-1D* plants exhibit a dwarf morphology.

SAR DEFICIENT 1 (SARD1) and CALMODULIN BINDING PROTEIN 60g (CBP60g) are two master transcription factors in plant immunity (Sun *et al*., 2015). Induction of SA biosynthetic genes such as *ISOCHORISMATE SYNTHASE 1 (ICS1), ENHANCED DISEASE SUSCEPTIBILITY 5 (EDS5)* and *AVRPPHB SUSCEPTIBLE 3 (PBS3)* during pathogen infection is coordinately regulated by SARD1 and CBP60g (Zhang *et al*., 2010a, Wang *et al*., 2011, Sun *et al*., 2015). Pathogen-induced SA accumulation is almost completely blocked in *sard1 cbp60g* double mutant plants (Zhang *et al*., 2010a, Wang *et al*., 2011). In addition, SARD1 and CBP60g are also required for activation of the biosynthesis of N-hydroxy-pipecolic acid (NHP), a signaling molecule important for both local resistance and SAR (Sun *et al*., 2020). They coordinately regulate pathogen-induced expression of NHP biosynthesis genes such as *AGD2-like defense response protein 1 (ALD1), SAR deficient 4 (SARD4)* and *Flavin-dependent monooxygenase 1 (FMO1)* (Sun *et al*., 2018, Sun *et al*., 2020). Both SARD1 and CBP60g belong to a small protein family with eight members in *Arabidopsis*. Unlike CBP60g and other members in the family which have calmodulin (CaM)-binding activity (Wang *et al*., 2009) (Reddy *et al*., 2002), SARD1 does not bind CaM and is primarily regulated at transcriptional level, as overexpression of *SARD1* is sufficient to activate downstream defense gene expression (Zhang *et al*., 2010a). Another member of the family, CBP60a, was shown to function as a negative regulator of plant immunity (Truman *et al*., 2013).

Both SARD1 and CBP60g contribute to autoimmunity of *snc2-1D* (Sun *et al*., 2015). Although the up-regulation of *ICS1* and accumulation of SA in *snc2-1D* are only slightly reduced in *cbp60g-1 snc2-1D* and *sard1-1 snc2-1D* double mutants, they are largely blocked in the *cbp60g-1 sard1-1 snc2-1D* triple mutant. The sizes of *cbp60g-1 snc2-1D* and *sard1-1 snc2-1D* double mutants are slight larger than *snc2-1D*, whereas the *cbp60g-1 sard1-1 snc2-1D* triple mutant exhibits wild type morphology. These findings suggest that there are two parallel pathways downstream of SNC2, one depending on SARD1 and the other requiring CBP60g (Sun *et al*., 2015). How the plasma membrane localized SNC2 transduces signals to SARD1 or CBP60g is largely unknown.

To identify components involved in the SARD1-dependent defense signaling downstream of SNC2, we performed a suppressor screen in the *cbp60g-1 snc2-1D* background. Here we report the identification and characterization of one of the suppressors, *bda7-1* (*bian da* 7; bian da means “becoming big” in Chinese). Positional cloning revealed that *BDA7* encodes another member of the CBP60 family, CBP60b, which positively regulates the expression of *SARD1*.

## Results

### Identification and characterization of *bda7-1 cbp60g-1 snc2-1D*

To identify regulators of SARD1-dependent defense signaling in SNC2-mediated immunity, we mutagenized *cbp60g-1 snc2-1D* and searched for mutants suppressing its dwarf morphology. *bda7-1* was one of the mutants identified. As shown in Figure 1A, *bda7-1 cbp60g-1 snc2-1D* has an intermediate size compared with wild type Col-0 and *cbp60g-1 snc2-1D*. Quantitative RT-PCR analysis showed that the elevated expression of *PR1* and *ICS1* in *cbp60g-1 snc2-1D* is fully blocked by *bda7-1* (Figure 1B and 1C). Consistent with the *ICS1* expression, both free (Figure 1D) and total SA (Figure 1E) levels in *bda7-1 cbp60g-1 snc2-1D* are considerably lower than in *cbp60g-1 snc2-1D*. In addition, the enhanced resistance against *Hpa* Noco2 observed in *cbp60g-1 snc2-1D* is lost in *bda7-1 cbp60g-1 snc2-1D* (Figure 1F). Together, these data indicate that *bda7-1* suppresses the autoimmunity of *cbp60g-1 snc2-1D*.

**Figure 1.**
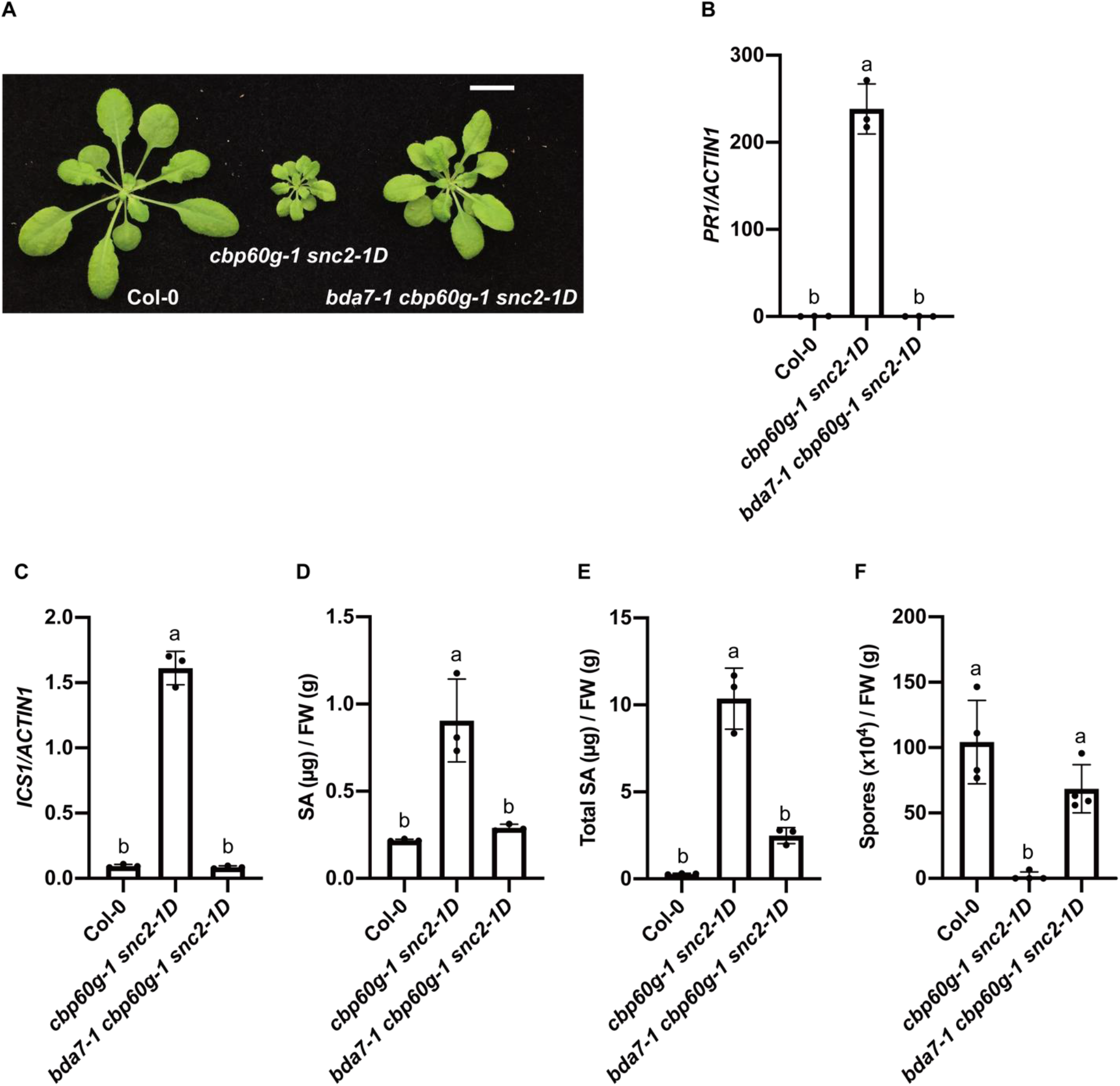
Identification and characterization of *bda7-1 cbp60g-1 snc2-1D*. (A) Morphology of four-week-old soil-grown plants of Col-0, *cbp60g-1 snc2-1D* and *bda7-1 cbp60g-1 snc2-1D* under long-day condition. Scale bar is 1 cm. (B and C) Relative expression levels of *PR1* (B) and *ICS1* (C) in the indicated genotypes. Transcript levels were normalized with *ACTIN1*. Error bars represent standard deviations. Letters indicate statistical differences (P < 0.0001, one-way ANOVA followed by Tukey’s multiple comparisons test; n = 3). (D, E) Free (D) and total SA (E) levels in the indicated genotypes. Error bars represent standard deviations. Letters indicate statistical differences (P < 0.01, one-way ANOVA followed by Tukey’s multiple comparisons test; n = 3). (F) Growth of *Hpa* Noco2 conidiospores on the indicated genotypes. Error bars represent standard deviations. Letters indicate statistical differences (P < 0.01, one-way ANOVA followed by Tukey’s multiple comparisons test; n = 4).

### Map-based cloning of *BDA 7*

When *bda7-1 cbp60g-1 snc2-1D* was backcrossed with *cbp60g-1 snc2-1D*, F1 plants showed similar morphology as *cbp60g-1 snc2-1D* (Figure S1), indicating that *bda7-1* is recessive. To map *bda7-1, bda7-1 cbp60g-1 snc2-1D* (in Col-0 background) was crossed with Landsberg *erecta* (L*er*). In the F2 population, plants homozygous for both *cbp60g-1* and *snc2-1D* with dwarf morphology were selected for linkage analysis. *bda7-1* was initially mapped to a region between SSLP markers MBG8 and MUB3 on chromosome 5, and subsequently fine-mapped to a ~0.8 Mb region between MUA2 and MMN10 (Figure S2).

To identify *BDA7*, DNA from 50 F2 plants with similar size as *bda7-1 cbp60g-1 snc2-1D* obtained from a cross between *bda7-1 cbp60g-1 snc2-1D* and *cbp60g-1 snc2-1D* were pooled and sequenced by Illumina sequencing. A single candidate gene within the ~0.8 Mb mapped region, *AT5G57580*, was found, which carries a homozygous C to T mutation in its coding sequence (Figure 2A and Figure S3), resulting in the substitution of Leu148 with Phe (Figure 2A).

**Figure 2.**
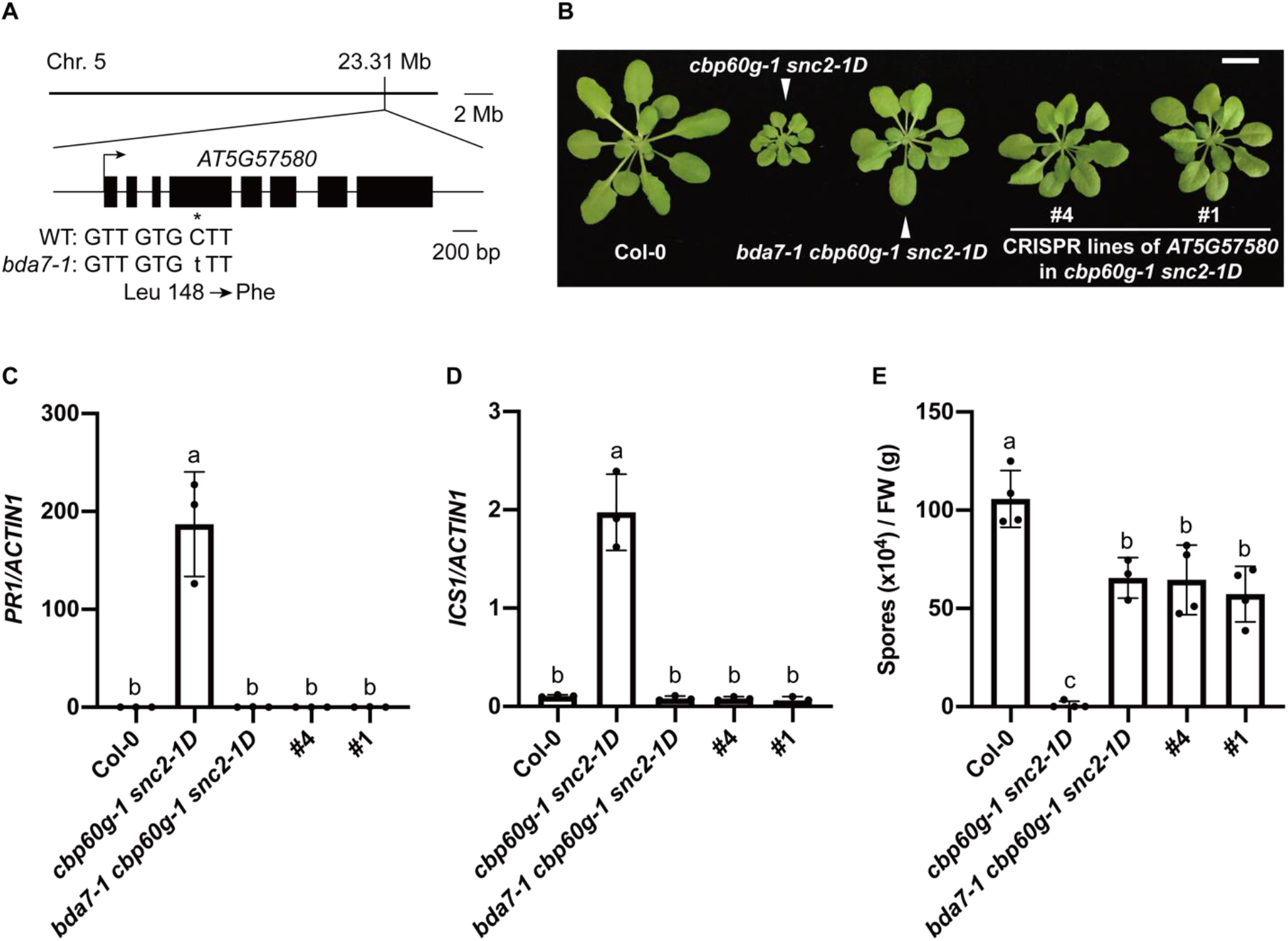
Deletion mutants of *AT5G57580* suppress the constitutively activated immunity of *cbp60g-1 snc2-1D*. (A) Map position and the mutation in *bda7-1*. The asterisk is used to mark the position of *bda7-1*. (B) Morphology of four-week-old soil-grown plants of the indicated genotypes under long-day condition. Scale bar is 1 cm. (C and D) Expression levels of *PR1* (C) and *ICS1* (D) in the indicated genotypes as normalized by *ACTIN1*. Error bars represent standard deviations. Letters indicate statistical differences (P < 0.01, one-way ANOVA followed by Tukey’s multiple comparisons test; n = 3). (E) Growth of *Hpa* Noco2 conidiospores on the indicated genotypes. Error bars represent standard deviations. Letters indicate statistical differences (P < 0.01, one-way ANOVA followed by Tukey’s multiple comparisons test; n = 4).

### Loss of *CBP60b* suppresses the autoimmunity of *cbp60g-1 snc2-1D*

To test whether *AT5G57580* is indeed required for the autoimmunity of *cbp60g-1 snc2-1D, AT5G57580* was knocked out by CRISPR/Cas9 in the *cbp60g-1 snc2-1D* background. As shown in Figure 2B, the newly generated *AT5G57580* CRISPR alleles suppressed *cbp60g-1 snc2-1D* similarly as *bda7-1*. Sanger sequencing confirmed that these lines carry homozygous deletion mutations in *BDA7* (Figure S4). Similar to *bda7-1*, these deletion mutants also suppress the constitutive expression of *PR1* and *ICS1* (Figure 2C and 2D), elevated SA levels (Figure S5) and enhanced resistance against *Hpa* Noco2 in *cbp60g-1 snc2-1D* (Figure 2E), suggesting that *AT5G57580* is required for the constitutively immune responses in *cbp60g-1 snc2-1D*.

To further confirm that suppression of the *cbp60g-1 snc2-1D* mutant phenotype in *bda7-1 cbp60g-1 snc2-1D* is caused by the mutation in *AT5G57580*, we transformed a construct expressing wild-type *AT5G57580* with a C-terminal 3×HA tag under the control of its native promoter into *bda7-1 cbp60g-1 snc2-1D*. As shown in Figure S6, the *AT5G57580-3HA* transgenic lines have similar morphology as *cbp60g-1 snc2-1D*, indicating that *AT5G57580-3HA* can complement the *bda7-1* mutation. Since *AT5G57580* encodes CBP60b, we renamed *bda7-1* as *cbp60b-1*.

The Leu148 residue in CBP60b is highly conserved among all CBP60 family members, except for CBP60a (Figure S7A), which functions as a negative regulator of plant immunity (Truman et al. 2013). Notably, from the same *cbp60g-1 snc2-1D* suppressor screen, three *sard1* alleles were found to carry mutations affecting Gly143, a residue close to the conserved Leu residue (Figure S7B). These data suggest that this region is important for the functions of CBP60 family proteins.

### CBP60b regulates *SARD1* expression

As the elevated *ICS1* expression and SA levels in *cbp60g-1 snc2-1D* are suppressed by *bda7-1* (Figure 1C-1E), we examined whether the expression of *SARD1* is affected in the *cbp60b cbp60g-1 snc2-1D* mutants. As shown in Figure 3A, the expression level of *SARD1* is much higher in *cbp60g-1 snc2-1D* than in wild type, and the elevated *SARD1* expression is fully suppressed in *cbp60b cbp60g-1 snc2-1D*, suggesting that CBP60b functions upstream of *SARD1*. We also examined the expression levels of *FMO1*, which is another target gene of SARD1. Consistent with the expression levels of *SARD1*, up-regulation of *FMO1* in *cbp60g-1 snc2-1D* is also blocked in *cbp60b cbp60g-1 snc2-1D* (Figure 3B). These data suggest that CBP60b functions upstream of SARD1.

**Figure 3.**
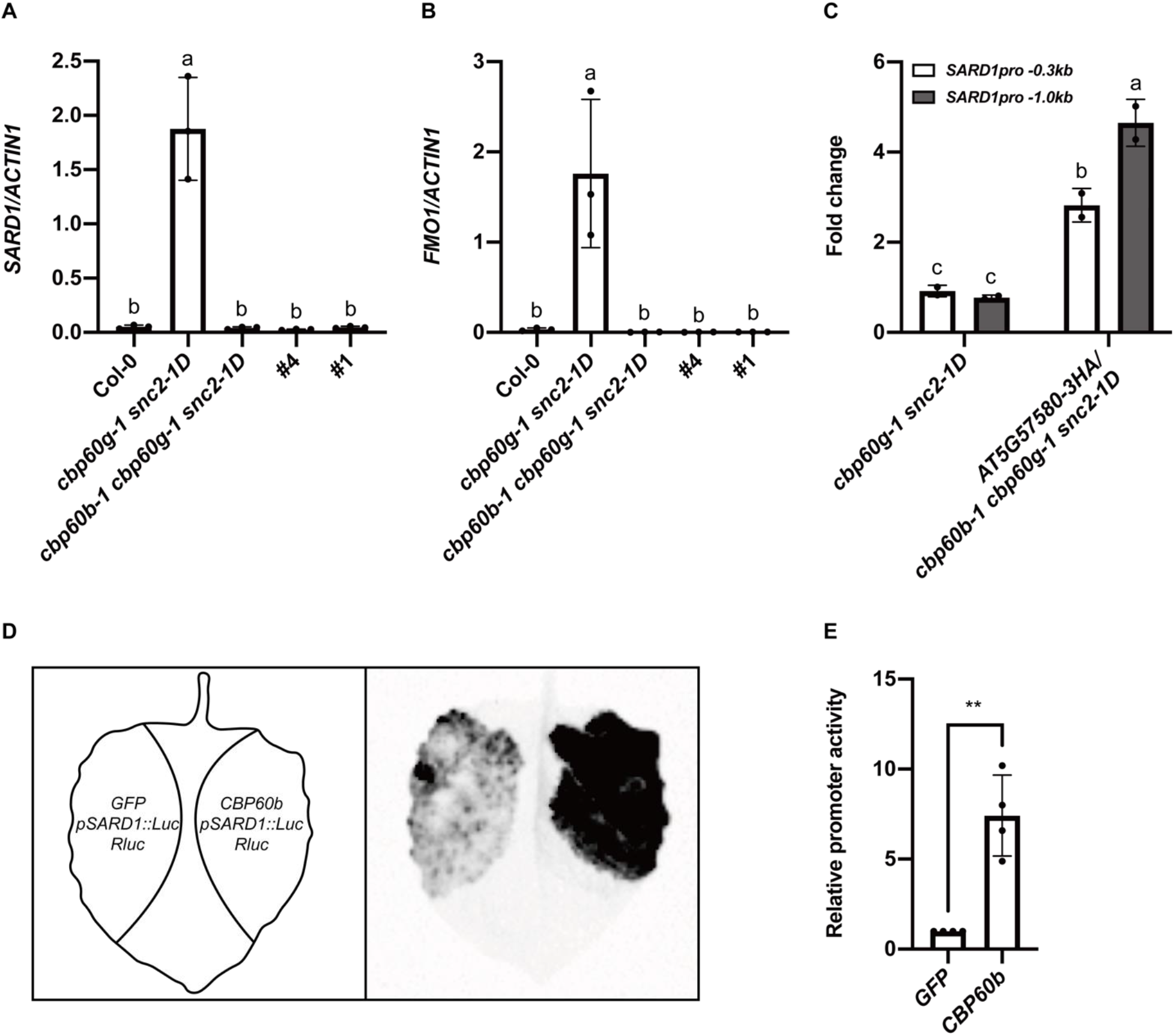
Regulation of *SARD1* expression by CBP60b. (A and B) Expression levels of *SARD1* (A) and *FMO1* (B) in the indicated genotypes as normalized by *ACTIN1*. Error bars represent standard deviations. Letters indicate statistical differences (P < 0.01, one-way ANOVA followed by Tukey’s multiple comparisons test; n = 3). (C) Binding of CBP60b to the promoter of *SARD1* as determined by ChIP-PCR. Two-week-old seedlings grown on ½ MS plates were sprayed with 1 μM of the elicitor nlp20 5 h before tissue collection. CBP60b-3HA chromatin complexes were immunoprecipitated with anti-HA antibody and protein G agarose beads. Negative control reactions were performed in parallel without adding anti-HA antibody. Immunoprecipitated DNA samples were quantified by qPCR using primers specific to *SARD1* promoter. ChIP results are presented as fold changes by dividing signals from ChIP with the anti-HA antibody by those from no antibody controls. Error bars represent standard deviations. Letters indicate statistical differences (P < 0.05, one-way ANOVA followed by Tukey’s multiple comparisons test; n = 2). (D) Activation of *pSARD1::Luc* reporter gene expression by CBP60b in *N. benthamiana* leaves. *Agrobacteria* carrying *pSARD1::Luc* (OD_600_=0.2) and *35S::Rluc* expressing the Renilla luciferase (OD_600_=0.05) were co-infiltrated with *Agrobacteria* carrying *35S::CBP60b* or *35S::GFP* (OD_600_=0.5). The left diagram illustrates the different treatments. 2 days after inoculation, luminescence was detected after infiltration with 1 mM luciferin, as shown in the right image. (E) Quantification of firefly luciferase activities in *N. benthamiana* leaves co-transformed with the construct combinations as indicated in (D). Relative promoter activity was presented as a ratio of firefly luciferase/renilla luciferase. Relative promoter activity of the GFP control group was set as 1. ** indicates statistical differences (P < 0.01, unpaired t test; n = 4).

As CBP60b is predicted to be a transcription factor, we tested whether CBP60b binds to the promoter of *SARD1*. ChIP-qPCR assay was carried out on transgenic plants expressing *CBP60b-3HA* under its own promoter in the *cbp60b-1 cbp60g-1 snc2-1D* background. Compared to the non-transgenic negative control *cbp60g-1 snc2-1D*, ~4.6-fold enrichment of the DNA around 1.0 kb upstream of the start codon and ~2.8 fold enrichment of the DNA around 0.3 kb upstream of the start codon were observed in the samples from the *CBP60b-3HA* transgenic line (Figure 3C), indicating that CBP60b is recruited to the *SARD1* promoter region.

To further test whether CBP60b can activate the expression of *SARD1*, a plasmid expressing a luciferase reporter gene under the control of the *SARD1* promoter (*pSARD1::Luc*) was transformed into *Nicotiana (N.) benthamiana*, together with a *35S::CBP60b-3HA* construct. Co-transformation of *35S::CBP60b-3HA* with *pSARD1::Luc* resulted in increased expression of the luciferase reporter (Figure 3D and 3E), suggesting that overexpression of *CBP60b* in *N. benthimiana* leads to activation of *SARD1* expression. Together, these data suggest that CBP60b serves as a transcriptional activator of *SARD1*.

### Overexpression of *CBP60b* leads to up-regulation of *SARD1* and enhanced disease resistance in *Arabidopsis*

To further test the hypothesis that CBP60b regulates *SARD1* expression, we overexpressed *CBP60b* in wild type Col-0 background. As shown in Figure 4A, transgenic plants overexpressing *CBP60b-3HA* exhibit dwarfism with dark green leaves. The expression levels of *SARD1* are significantly higher in the *CBP60b-3HA* transgenic lines than in the wild type control (Figure 4B), further supporting the role of CBP60b in regulating *SARD1* expression. Consistent with the elevated *SARD1* expression levels, these transgenic lines showed enhanced resistance to *Hpa* Noco2 (Figure 4C).

**Figure 4.**
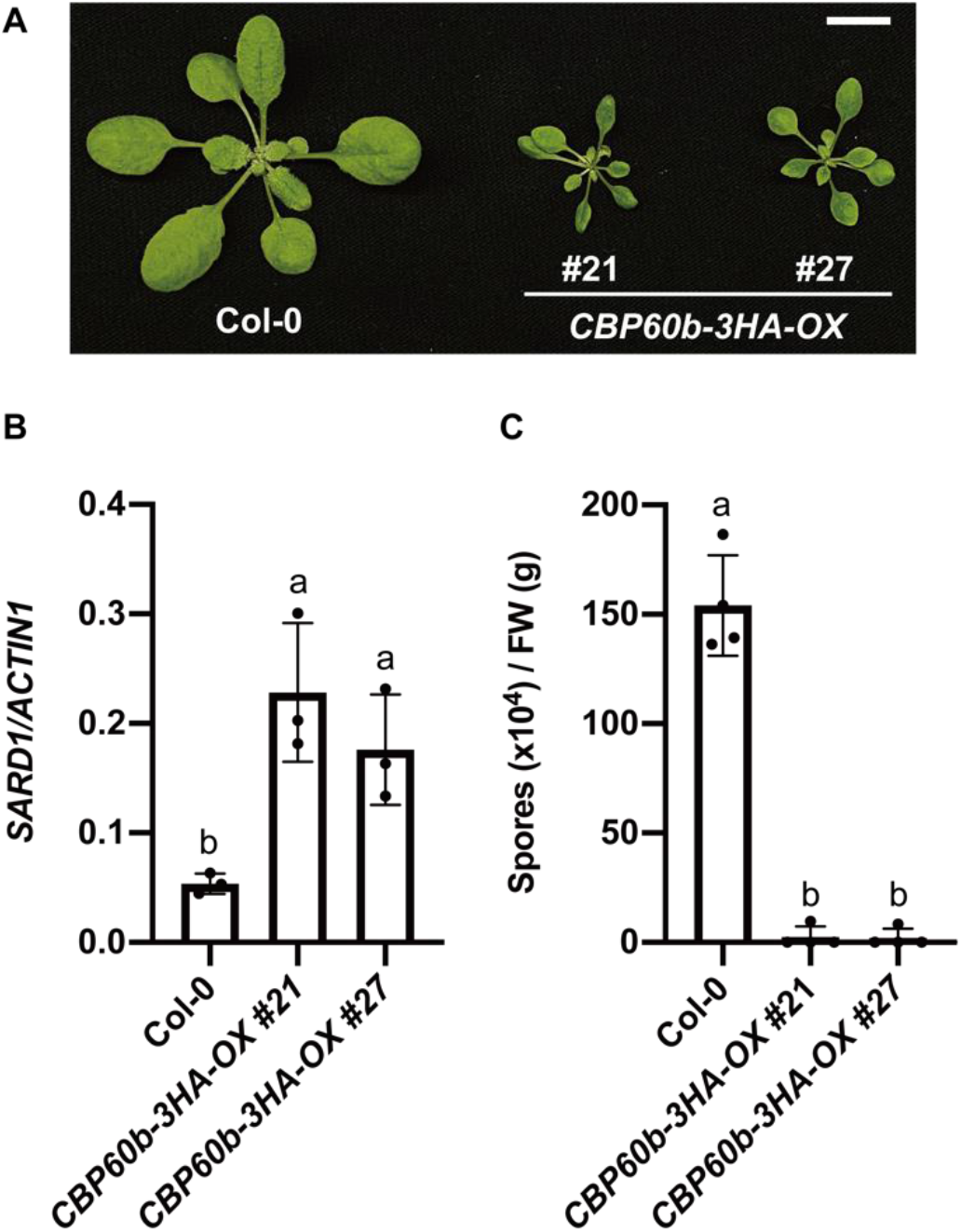
Overexpression of *CBP60b* leads to increased *SARD1* expression and enhanced resistance against *Hpa* Noco2. (A) Morphologies of three-week-old soil-grown plants of the indicated genotypes under long-day condition. Scale bar is 1 cm. (B) Expression level of *SARD1* in the indicated genotypes as normalized by *ACTIN1*. Error bars represent standard deviations. Letters indicate statistical differences (P < 0.05, one-way ANOVA followed by Tukey’s multiple comparisons test; n = 3). (C) Growth of *Hpa* Noco2 on the indicated genotypes. Error bars represent standard deviations. Letters indicate statistical differences (P < 0.0001, one-way ANOVA followed by Tukey’s multiple comparisons test; n = 4).

### *cbp60b* single mutants exhibit constitutively activated immune responses

The requirement of CBP60b for the constitutive defense responses in *cbp60g-1 snc2-1D* indicates that CBP60b serves as a positive regulator of plant immunity. To our surprise, when we isolated the *cbp60b-1* single mutant from the F2 population of a cross between *cbp60b-1 cbp60g-1 snc2-1D* and Col-0, the *cbp60b-1* plants displayed a dwarf morphology with dark green and abnormal leaves (Figure S8). *cbp60b-1 cbp60g-1* double mutant plants isolated from the same F2 population showed an even more dramatic phenotype, as they were seedling-lethal and grew only two cotyledons at room temperature (Figure S8). To rule out the possibility that the unexpected phenotypes originated from random background mutations in *cbp60b-1 cbp60g-1 snc2-1D*, we generated *cbp60b* deletion mutants in wild type Col-0 and *cbp60g-1* backgrounds by CRISPR/Cas9. As shown in Figure 5A, the *cbp60b* deletion mutants showed almost identical morphology as the corresponding *cbp60b-1* mutants in wild type and *cbp60g-1* backgrounds, confirming that the dwarfism observed in *cbp60b-1* and *cbp60b-1 cbp60g-1* is caused by the mutation in *CBP60b*.

**Figure 5.**
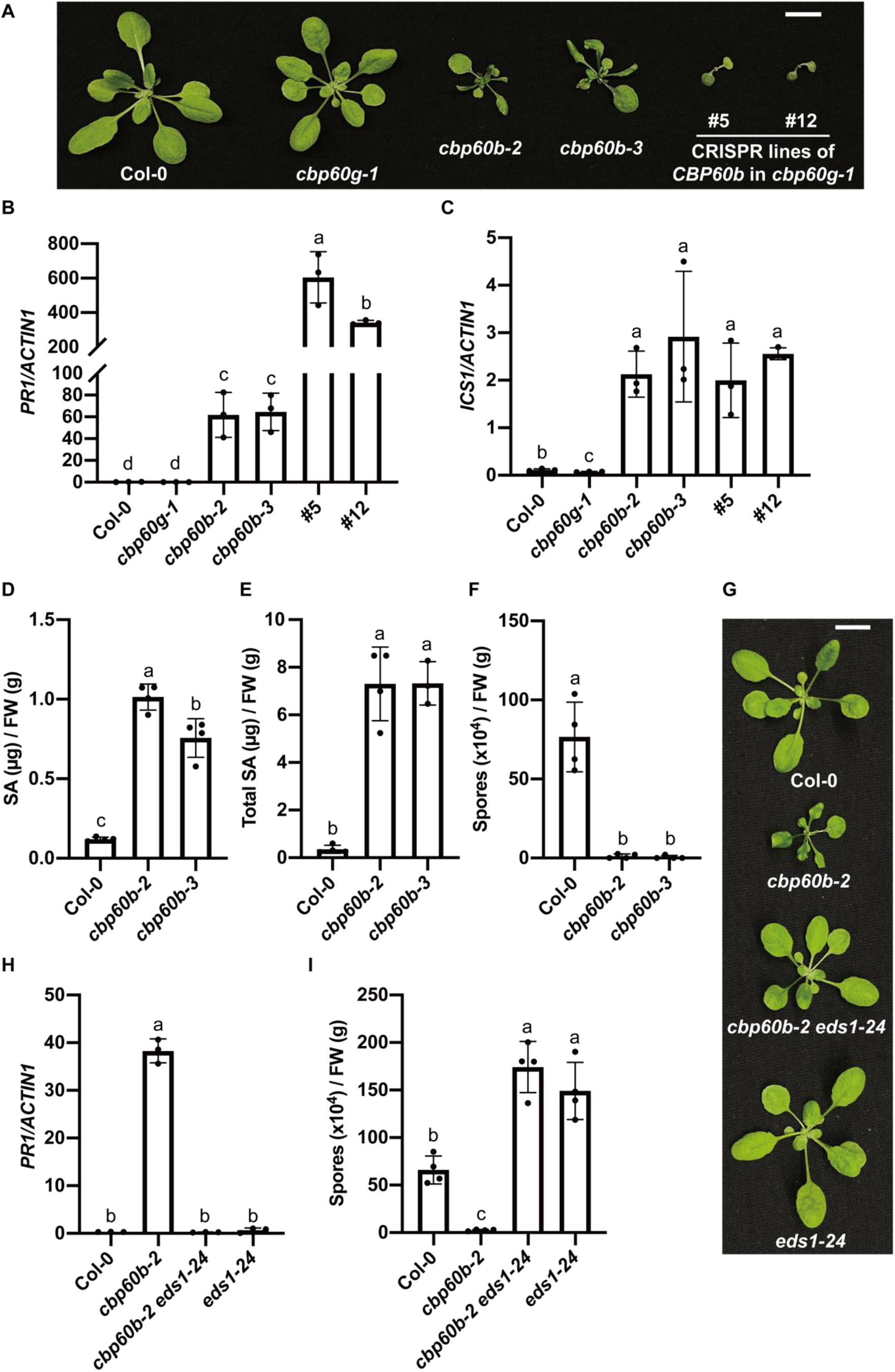
Knocking out *CBP60b* activates *EDS1*-dependent defense responses. (A, G) Morphologies of three-week-old soil-grown plants of the indicated genotypes under long-day condition. Scale bar is 1 cm. (B, C) Expression levels of *PR1* (B) and *ICS1(C)* in the indicated genotypes as normalized by those of *ACTIN1*. Error bars represent standard deviations. Letters indicate statistical differences (P < 0.05, Student’s t-test; n = 3). (D, E) Free (D) and total SA(E) levels in the indicated genotypes. Error bars represent standard deviations. Letters indicate statistical differences (P < 0.01, one-way ANOVA followed by Tukey’s multiple comparisons test; n = 4). (F) Growth of *Hpa* Noco2 on the indicated genotypes. Error bars represent standard deviations. Letters indicate statistical differences (P < 0.0001, one-way ANOVA followed by Tukey’s multiple comparisons test; n = 4). (H) Expression levels of *PR1* in the indicated genotypes as normalized by those of *ACTIN1*. Error bars represent standard deviations. Letters indicate statistical differences (P < 0.0001, one-way ANOVA followed by Tukey’s multiple comparisons test; n = 3). (I) Growth of *Hpa* Noco2 on the indicated genotypes. Error bars represent standard deviations. Letters indicate statistical differences (P < 0.01, one-way ANOVA followed by Tukey’s multiple comparisons test; n = 4).

Next, we tested whether defense responses are activated in these mutants. As shown in Figure 5B, the expression levels of *PR1* were dramatically increased in *cbp60b* mutants, and even higher in the *cbp60b cbp60g-1* double mutants. *cbp60b* and *cbp60b cbp60g-1* mutants also exhibited elevated *ICS1* expression (Figure 5C). Consistently, SA levels in *cbp60b* mutants were much higher than in the wild type (Figure 5D and 5E). In addition, *cbp60b* mutants showed strong resistance against *Hpa* Noco2 (Figure 5F). These observations suggest that knocking out *CBP60b* leads to constitutive activation of defense responses.

We further examined whether mutations in *SARD1* affects the autoimmunity of *cbp60b cbp60g-1. cbp60b cbp60g-1 sard1-1* triple mutants were generated through knocking out *CBP60b* in the *cbp60g-1 sard1-1* double mutant by CRISPR/Cas9. Similar to the *cbp60b cbp60g-1* double mutant, *cbp60b cbp60g-1 sard1-1* triple mutants were seedling lethal and grew only two cotyledons (Figure S9A). Unlike *cbp60b cbp60g-1*, cotyledons of *cbp60b cbp60g-1 sard1-1* plants also showed visible lesion (Figure S9A).

Positive regulators of plant immunity are often monitored/guarded by NLR receptors (Cui *et al*., 2015). To test whether the constitutive defense responses in *cbp60b* is caused by activation of NLR-mediated immunity, we crossed *cbp60b-2* with *eds1-24*, a CRISPR deletion knockout mutant of *EDS1*, a gene known to be required for defense signaling mediated by Toll/interleukin-1 receptor (TIR) domain containing NLRs (TNLs)(Aarts *et al*., 1998). The *cbp60b-2 eds1-24* double mutant showed wild type-like morphology (Figure 5G). In addition, the constitutive *PR1* expression and enhanced resistance to *Hpa* Noco2 are completely suppressed in *cbp60b-2 eds1-24* (Figure 5H and 5I), indicating that the autoimmunity of *cbp60b* is dependent on *EDS1*, and TNL-mediated immunity is likely activated with loss of *CBP60b* function.

As NLR-mediated autoimmunity can often be suppressed by high temperature (van Wersch *et al*., 2016), we tested whether the dwarf phenotype of *cbp60b, cbp60b cbp60g-1* and *cbp60b cbp60g-1 sard1-1* can be suppressed by growing them at 28 °C. As shown in Figure S9B, *cbp60b* exhibited wild type morphology at 28 °C, while *cbp60b cbp60g-1* and *cbp60b cbp60g-1 sard1-1* plants had intermediate size and were able to grow true leaves at 28 °C.

## Discussion

SARD1 and CBP60g play critical roles in transcriptional regulation of plant immunity (Sun *et al*., 2015). Unlike CBP60g, which requires binding of CaM for its activation (Wang *et al*., 2009), SARD1 is primarily regulated at transcription level. The autoimmune mutant *snc2-1D* provides a unique system to uncover components regulating *SARD1*expression, as the constitutive defense responses in *cbp60g-1 snc2-1D* is mainly dependent on *SARD1* (Sun *et al*., 2015). From a suppressor screen of *cbp60g-1 snc2-1D*, here we identified CBP60b as a positive regulator of *SARD1* transcription. CBP60b is targeted to the promoter region of *SARD1* and required for its up-regulation in *cbp60g-1 snc2-1D*. Overexpression of *CBP60b* leads to elevated *SARD1* expression and constitutive defense responses. In addition, transient expression of *CBP60b* in *N. benthamiana* activates the expression of the *pSARD1-Luc* reporter gene, confirming that CBP60b positively regulates the expression of *SARD1*.

CBP60b belongs to the same protein family as SARD1 and CBP60g (Reddy *et al*., 2002), which share a highly conserved central domain with DNA-binding activity but have divergent sequences at the N- and C-termini (Wang *et al*., 2009, Zhang *et al*., 2010a). Unlike *SARD1* and *CBP60g*, which are strongly induced during pathogen infection (Wang *et al*., 2009, Zhang *et al*., 2010a), *CBP60b* is constitutive expressed in different tissue (Reddy *et al*., 2002). CBP60b was identified as a CaM-binding protein, suggesting that its activity might be influenced by Ca^2+^ levels (Reddy *et al*., 2002). SARD1 was previously shown to activate *ICS1* expression through a

GAAATTT motif on its promoter (Sun et al. 2015). CBP60b shares ~80% sequence similarity with SARD1 in the middle domain. Whether it binds to a similar DNA sequence in activating *SARD1* expression remains to be determined.

Surprisingly, defense responses are constitutively activated in the *cbp60b* single mutants. The constitutive defense response in *cbp60b* is blocked by knocking out *EDS1*, suggesting that loss of *CBP60b* leads to activation of TNL-mediated immune signaling. It is possible that CBP60b is required for the expression of a negative regulator of TNL-mediated immune signaling. More likely, CBP60b may be monitored by a TNL and loss of its function triggers activation of this unidentified TNL and downstream defense responses, since critical immune regulators are often targeted by pathogen effector proteins and guarded by plant NLR receptors (Cui *et al*., 2015, Kourelis and Van Der Hoorn, 2018). Although no pathogen effector has been reported to target CBP60b, SARD1 and CBP60g were shown to be targets of *Verticillium* effector VdSCP4 (Qin *et al*., 2018).

Compared to *cbp60b*, the autoimmune phenotype of the *cbp60b cbp60g-1* double mutants is even more dramatic, suggesting that *CBP60b* and *CBP60g* play partially redundant roles in transcriptional regulation of the putative guardee/decoy recognized by the unknown TNL. In *cbp60b*, the expression of the putative guardee/decoy is partially blocked, leading to activation of TNL-mediated immunity. Lacking both CBP60b and CBP60g leads to further reduction of its expression and stronger defense responses. Interestingly, the severe dwarf phenotype of *cbp60b cbp60g-1* was not observed in the *snc2-1D* background. It is possible that *snc2-1D* can also activate CBP60b and CBP60g-inpendent expression of the putative guardee/decoy, which compensates the loss of CBP60b and CBP60g.

A working model is proposed based on our findings (Figure 6). CBP60b promotes the expression of *SARD1* and contributes to SNC2-mediated immunity. It is required for the expression of a yet-to-be-identified guardee/decoy of a TNL. Disruption of the expression of the putative guardee/decoy results in activation of TNL-mediated immunity. It is of future interest to identify this guardee/decoy and the corresponding TNL.

**Figure 6.**
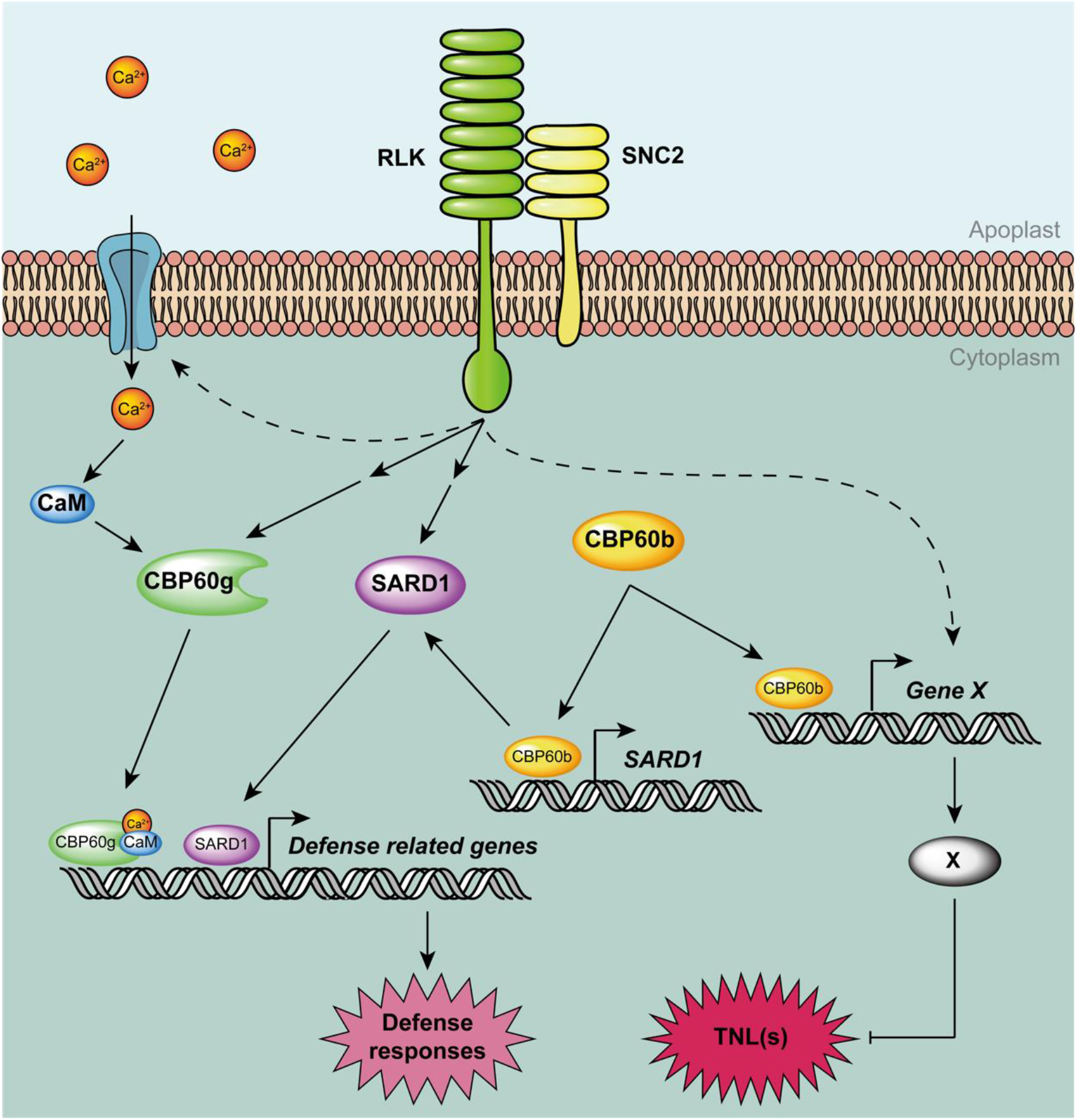
A working model of CBP60b in plant immunity. On one hand, CBP60b contributes to SNC2-mediated immunity by promoting *SARD1* expression. On the other hand, it is also required for the expression of an unknown gene (X), which encodes a guardee/decoy of a TNL(s) or a negative regulator of TIR signaling. Activation of SNC2-mediated defense responses also leads to CBP60b-independent expression of gene X.

## Materials and Methods

### Plant materials and growth conditions

All *Arabidopsis thaliana* mutants are in the Col-0 ecotype background unless specified. The *cbp60g-1 snc2-1D, cbp60g-1, cbp60g-1 sard1-1* plants were described previously (Zhang *et al*., 2010a, Sun *et al*., 2015). *cbp60b-1* and *cbp60b-1 cbp60g-1* plants were isolated from a cross between *cbp60b-1 cbp60g-1 snc2-1D* and Col-0. *cbp60b-2, cbp60b-3* are two independent deletion lines generated by CRISPR/Cas9. The *CBP60b-3HA-OX #21* and *#27* are two independent overexpression lines generated by transforming Col-0 with *Agrobacteria* carrying pCambia1300-35S-CBP60b-3HA. *eds1-24* is a CRISPR mutant line with deleting both *EDS1A* and *EDS1B* genes, as described previously (Tian *et al*., 2020). *cbp60b-2 eds1-24* was obtained from the F2 population of a cross between *cbp60b-2* and *eds1-24*. Primers used for genotyping were listed in Table S1.

Plants were grown on soil under long-day conditions (16-h light/ 8-h dark cycle) with a light intensity of ~100 μmol/m^2^/s of at 22 °C unless specified. Plants for quantitative RT-PCR were grown on plates with ½ Murashige and Skoog (MS) and 1% sucrose for two weeks. Plants used for SA quantification were grown on soil for four weeks under short-day conditions (8-h light/ 16-h dark cycle).

### Constructs for generation of deletion mutants and transgenic plants

The CRISPR/Cas9 system used for generating *cbp60b* mutants was described previously (Xing *et al*., 2014). Two guide RNAs were designed to target *CBP60b* genomic DNA for generation of a deletion of ~1 kb in size. A PCR fragment containing the guide RNA sequences was amplified from the pCBC-DT1T2 vector using primers At5g57580-BsFF0 and At5g57580-BsRR0 and subsequently inserted into the pHEE401 vector using the BsaI site. The derived plasmid was transformed into *E. coli* DH10B and later *Agrobacterium* GV3101 by electroporation. Arabidopsis plants were transformed with the *Agrobacterium* carrying the plasmid by floral dipping (Clough and Bent, 1998). T1 plants were analyzed for deletion in *CBP60b* by PCR with primers listed in Table S1. Homozygous deletion mutants were obtained in the T2 generation.

To overexpress *CBP60b, CBP60b* was amplified from genomic DNA using primers AT5G57580-atgKpnI-F and AT5G57580-nstopBamHI-R. The PCR fragment was digested with KpnI and BamHI and afterwards ligated into pCambia1300-35S-3HA vector. The derived plasmid was transformed into *E. coli* DH10B and later *Agrobacterium* GV3101. Wild type Col-0 plants were transformed with *Agrobacterium* carrying the plasmid by floral dipping. Homozygous transgenic lines were obtained in the T2 generation.

For transgene complementation, a genomic DNA fragment containing *CBP60b* was amplified with primers AT5G57580-KpnI-F and AT5G57580-nstopBamHI-R. The PCR fragment was inserted into a pBasta-3HA vector derived from pCambia1305 to obtain the pAt5g57580::At5g57580-3HA construct. The *pAt5g57580::At5g57580-3HA/cbp60b-1 cbp60g-1 snc2-1D* transgenic lines were obtained by transforming *cbp60b-1 cbp60g-1 snc2-1D* plants with *Agrobacterium* carrying the plasmid of *pAt5g57580::At5g57580-3HA*.

### Chromatin immunoprecipitation (ChIP) analysis

ChIP-qPCR assays were performed as previously described (Sun *et al*., 2015). The chromatin complexes containing CBP60b-3HA were immunoprecipitated using anti-HA antibody (Roche, Basel, Switzerland) and Protein A/G Agarose beads (GE Healthcare, Chicago, United States). The immunoprecipitated DNA was analyzed by qPCR using gene specific primers, which were listed in Table S1.

### Genetic mapping and identification of *bda7-1*

Genetic mapping of *bda7-1* was carried out on the F2 population of a cross between *bda7-1 cbp60g-1 snc2-1D* mutant (in Col-0 background) and Landsberg *erecta* (L*er*). Afterwards, Illumina sequencing was used to identify the *bda7-1*mutation. *bda7-1 cbp60g-1 snc2-1D* was backcrossed with *cbp60g-1 snc2-1D*. Genomic DNA from 50 F2 plants with similar morphology as *bda7-1 cbp60g-1 snc2-1D* were pooled and sequenced.

### Dual reporter assay

Dual reporter assay was performed in *Nicotiana benthamiana* by transforming the reporter constructs together with the different effector constructs. The *pSARD1::Luc* were described previously (Ding *et al*., 2018). A *pUBQ1-driven* Renilla luciferase reporter was included as internal control. *Agrobacteria* carrying the reporter construct *pSARD1::Luc*, internal control *pUBQ1-Renilla* and effector construct *35S::CBP60b-3HA* or *35S::GFP-3HA* were first cultured in liquid LB and then resuspended in 10mM MgCl_2_. Leaves of *N. benthamiana* were co-infiltrated with *Agrobacteria* carrying the indicated construct combinations with final concentrations of OD_600_ = 0.2 (*pSARD1::Luc*), OD_600_ = 0.05 (*pUBQ1-Renilla*) and OD_600_ = 0.5 (*35S::CBP60b-3HA* or *35S::GFP-3HA*). For imaging the luminescence intensity, *N. benthamiana* leaves was infiltrated with 1 mM luciferin 40 h after inoculation of the *Agrobacteria* and then imaged under the Gel Doc XR+ System (Bio-Rad) with the Bolt mode. For quantification of the promoter activity, areas of *N. benthamiana* leaves inoculated with *Agrobacteria* was collected 40 h after inoculation. The Dual-Luciferase^®^ Reporter Assay System (Promega) was used to measure the activity of firefly luciferase and renilla luciferase sequentially using a BioTek™ Synergy™ 2 Multi-Mode Microplate Reader. Relative promoter activities were calculated as the ratio of firefly luciferase/renilla luciferase.

### RNA extraction, reverse transcription and qPCR

Plants for gene expression assays were grown on ½ MS plates for 14 days under long-day conditions. Approximately 50 mg plant tissue from 3-4 individual seedlings of the indicated genotypes were collected as a single sample. Three biological replicates were analyzed for each genotype. RNA extraction was performed using the EZ-10 Spin Column Plant RNA Miniprep Kit (Bio Basic Inc., Toronto, Canada). RNAs were reverse transcribed into cDNAs by OneScript Reverse Transcriptase (Applied Biological Materials Inc., Richmond, Canada). qPCR was performed on the total cDNAs using SYBR Premix Ex TaqTM II (Takara, Shiga, Japan). Primers for qPCR are listed in Table S1.

### SA extraction and quantification

The procedure of SA extraction and measurement was reported previously (Sun *et al*., 2015). ~100 mg of plant tissue was collected from 2-3 individual plants of the indicated genotypes as a single sample. Plant tissue was ground into fine powder with liquid nitrogen, resuspended with 600 μl 90% methanol and sonicated for 20 min to release SA. After centrifugation at 12,000×g for 10 min, the supernatant was collected, and the pellets underwent a second round of extraction by adding 500 μl of 100% methanol and sonicating for another 20 min. The supernatant from both extractions were combined together and dried by vacuum. Next, 500 μl of 5% (w/v) trichloroacetic acid was added to the dry samples, vortexed and sonicated for 5 min. After centrifugation at 12,000g for 15 min, the supernatant was collected and extracted with 500 μl extraction buffer (ethylacetate acid: cyclopentane: isoporopanal, 100:99:1, v/v/v) for three times. Each time, after centrifugation at 12,000×g for 10 min, the organic phase was collected and combined into a new tube. The combined organic phase was then dried by vacuum. The dry sample was next resuspended with 200 μl mobile phase (0.2M KAc, 0.5mM EDTA pH=5) by vortexing and sonicating for 5 min. After the final centrifugation at 12,000×g for 5 min, the supernatant was collected and used to determine the amount of SA by high-performance liquid chromatography.

### Pathogen infection assay

*Hpa* Noco2 infection was conducted by spray-inoculating two-week-old seedlings with spores in water (50,000 spores/mL). Inoculated seedlings were covered with a transparent lid and grown in a plant chamber at 18 °C with a relative humidity of ~80%. Infection was scored at 7 dpi by counting conidia spores with a hemocytometer. 4-5 individual plants were pooled as a single sample. 4 biological replicates were included for each genotype.

### Statistical analysis

Error bars in all of the figures represent standard deviations. The number of biological replicates is indicated in the figure legends. Statistical comparison among different samples is carried out by either one-way ANOVA with Tukey’s honestly significant difference (HSD) post hoc test or Student’s t-test, as reported in the figure legends.

## Acknowledgement

We would like to acknowledge Dr. Tongjun Sun for suggestions on the ChIP assay. We are grateful to Sean Shang for his help with the NGS data pipeline. We thank Jia Yao for her technical supports on the project. We thank Dr. Yanan Liu, Dr. Xingchuan Huang and Dr. Wei Li for the help with handling the sequencing samples.

## Author contributions

YZ and XL designed and supervised the study. WH, WZ and HT conducted the experimental research. WH, XL and YZ analyzed the data and wrote the manuscript.

