## Supporting information for "Arabidopsis CALMODULIN-BINDING PROTEIN 60b plays dual roles in plant immunity"

**Supplementary figures**

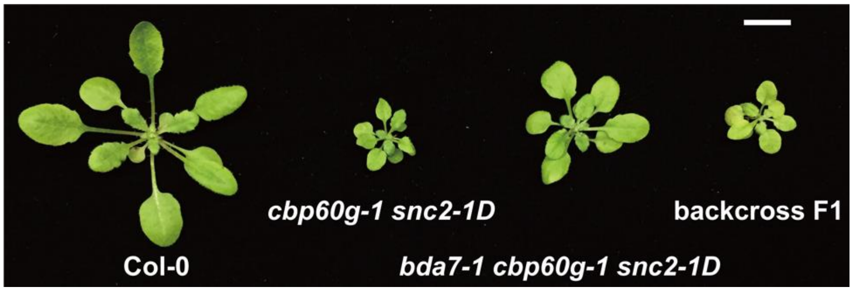


**Figure S1** *bda7-1* is recessive. Morphologies of three-week-old soil-grown plants of Col-0, *cbp60g-1 snc2-1D*, *bda7-1 cbp60g-1 snc2-1D* and the backcrossed F1 plant under long-day condition. Scale bar is 1 cm.

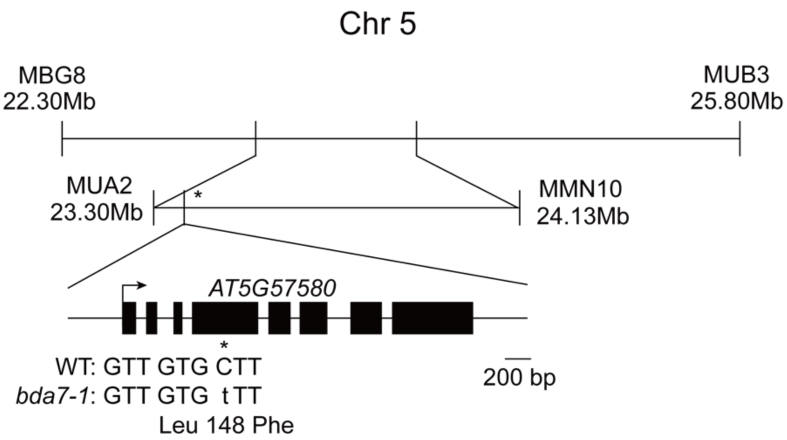


**Figure S2** Map position and the mutation in *bda7-1*. The asterisk is used to mark the position of *bda7-1*.

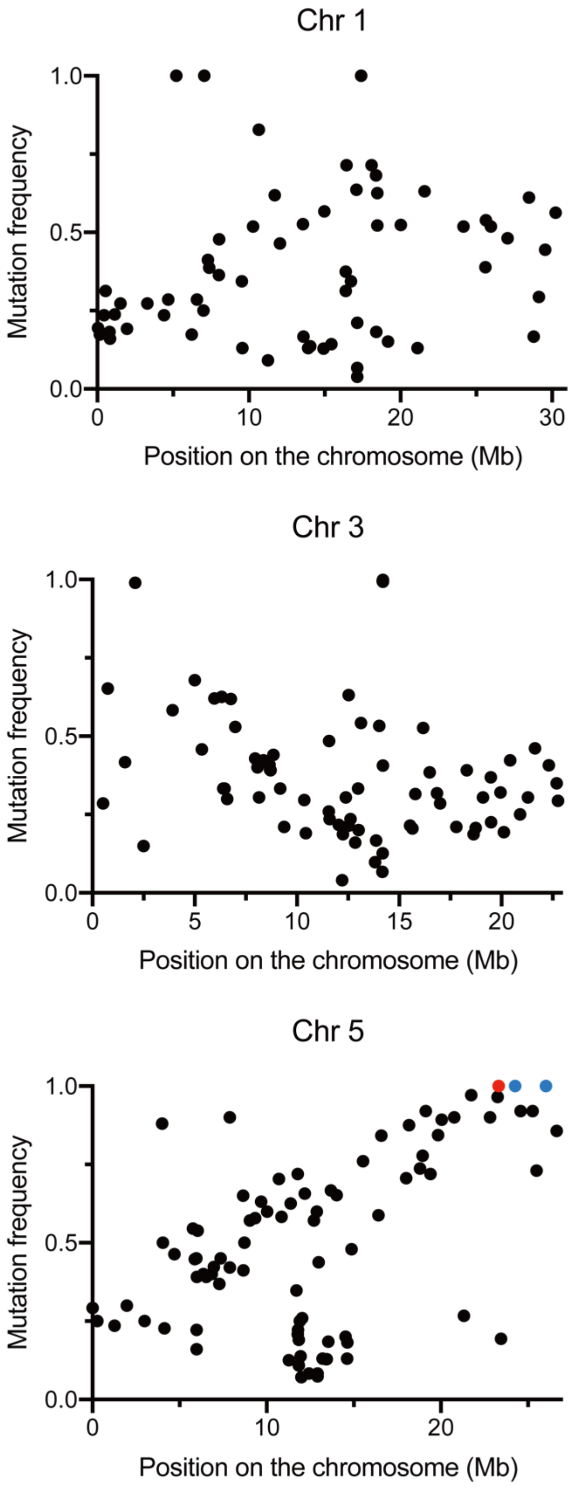


**Figure S3** Dot plots of frequency (*f*) of SNP mutations on five chromosomes in *bda7-1 cbp60g-1 snc2-1D*, as derived from Illumina sequencing. Colored dots represent mutations with a *f*=1. The red dot indicates the mutation with a *f*=1 within the mapped region.

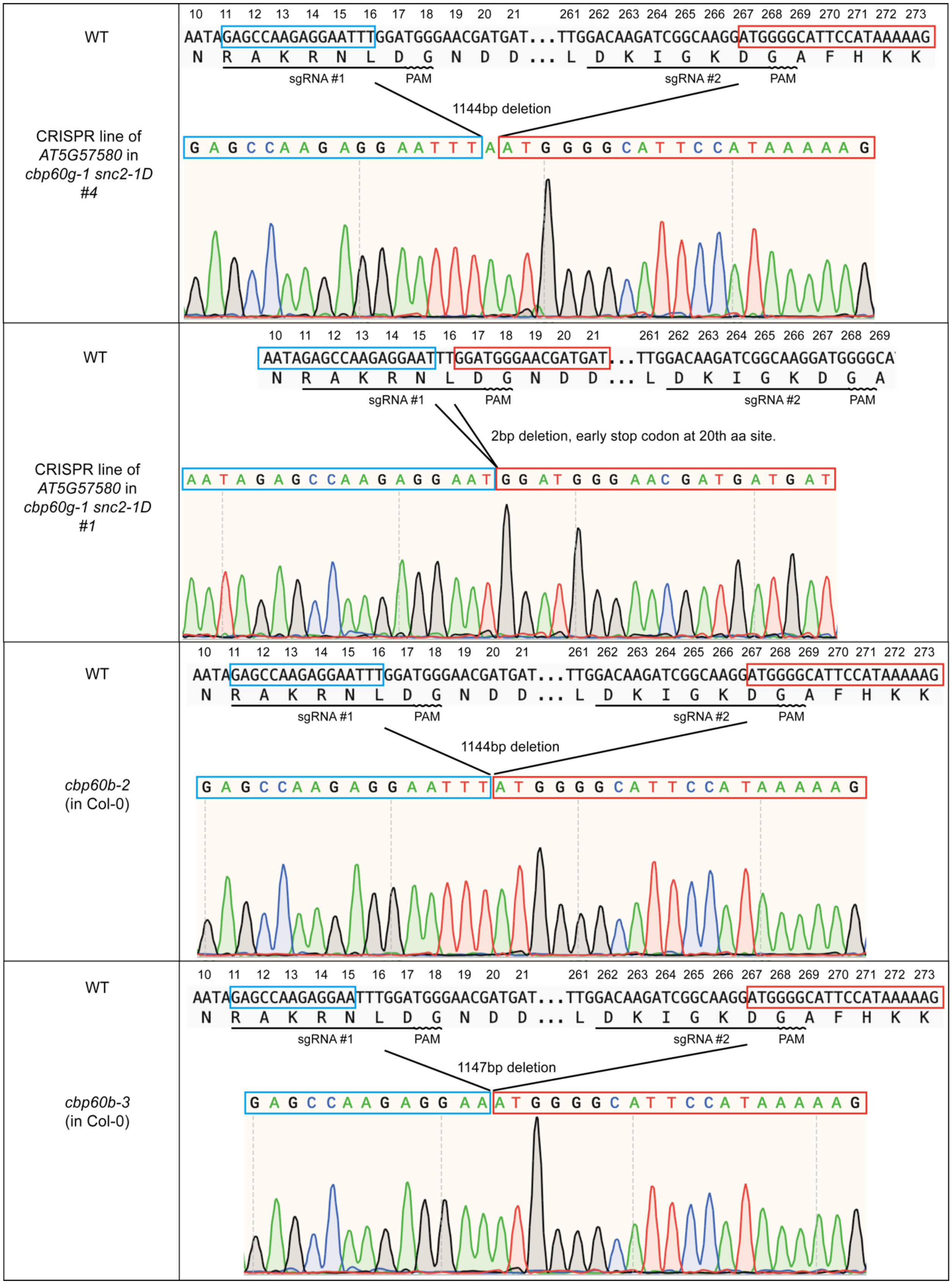


**Figure S4** Sequences of deletion mutations in *CBP60b* generated by CRISPR/Cas9.

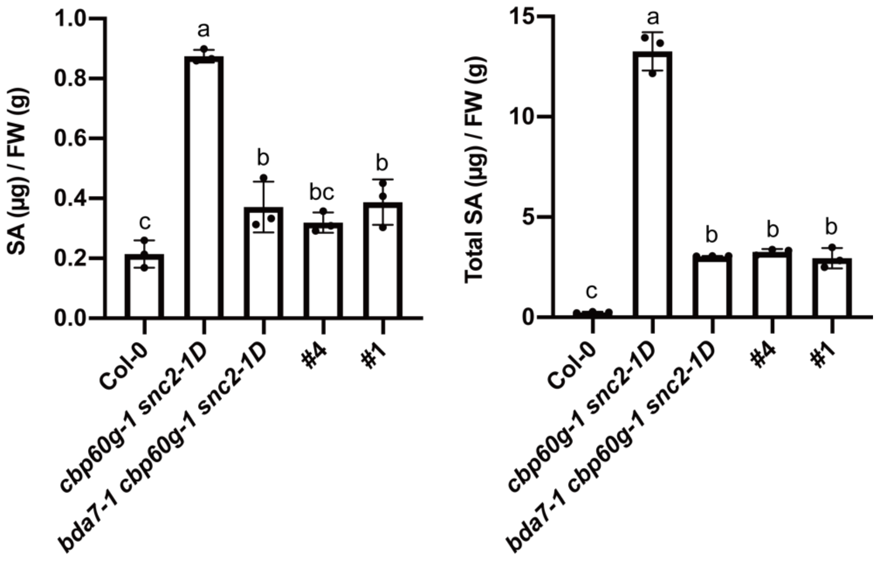


**Figure S5** Free and total SA levels in wild type (Col-0), *cbp60g-1 snc2-1D*, *bda7-1 cbp60g-1 snc2-1D*, and two *cbp60b* CRISPR-Cas9 deletion lines (#1 and #4) in *cbp60g-1 snc2-1D* background. Error bars represent standard deviations. Letters indicate statistical differences (P < 0.05, one-way ANOVA followed by Tukey's multiple comparisons test; n = 3).

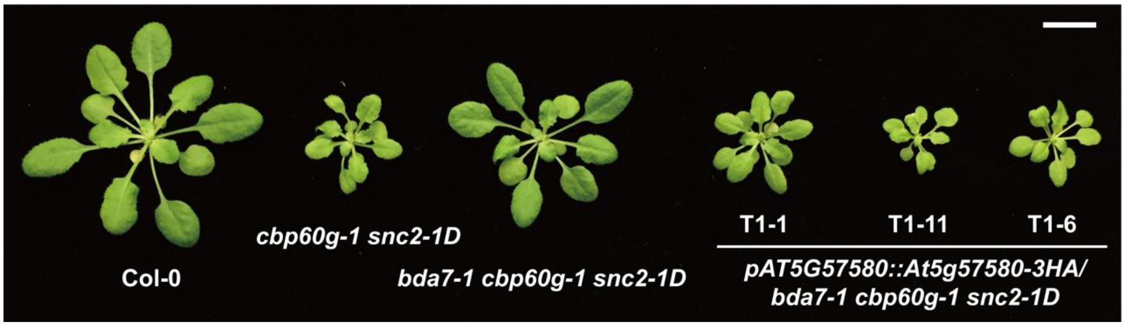


**Figure S6** *bda7-1* can be complemented by *AT5G57580-3HA* driven by its native promoter. Morphologies of four-week-old soil-grown plants of the indicated genotypes under long-day condition. Scale bar is 1 cm.

A

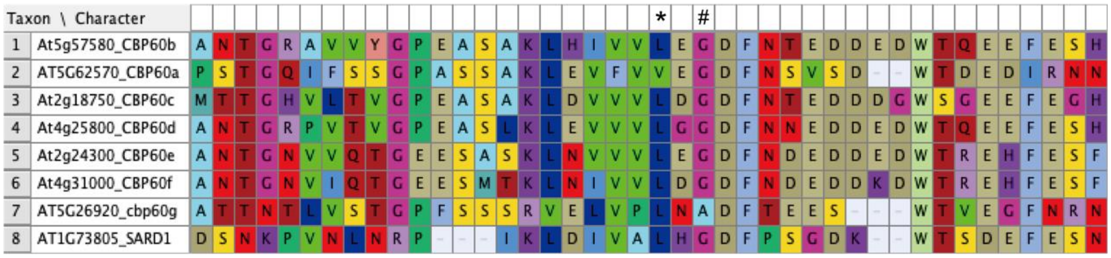


B

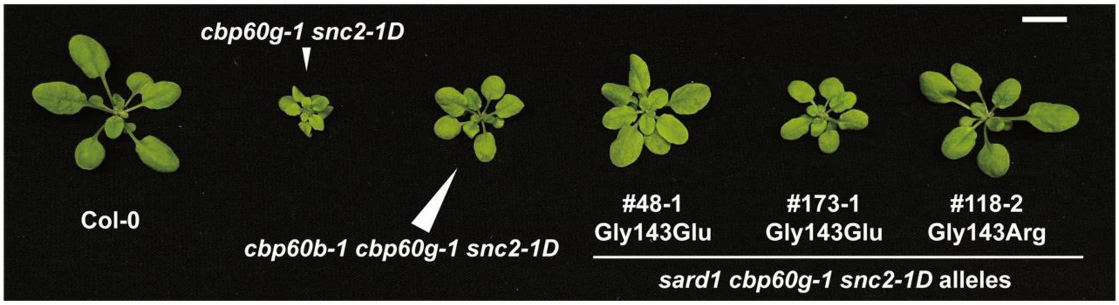


**Figure S7** The region containing Leu148 is important for the function of CBP60b and SARD1. (A) Alignment of protein sequences of CBP60 family members showing the conserved region containing the Leu148 residues. The * marks the Leu148 site in CBP60b. The # marks the Gly143 site in SARD1. Full-length protein sequences were aligned using MUSCLE (Edgar 2004). The alignment was presented by Mesquite (Maddison 2019).

(B) Morphologies of three-week-old soil-grown plants of the indicated genotypes under long-day condition. Scale bar is 1 cm.

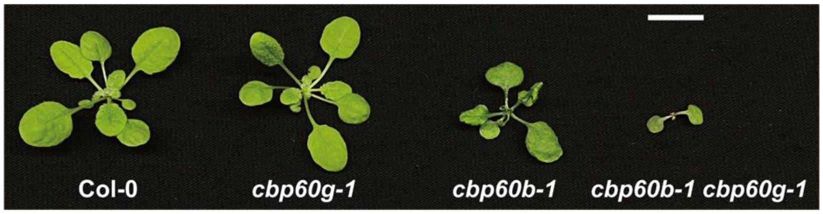


**Figure S8** The morphology of *cbp60b-1* single and *cbp60b-1 cbp60g-1* double mutant plants. The photo was taken on three-week-old soil-grown plants of the indicated genotypes under long-day condition. Scale bar is 1 cm.

A

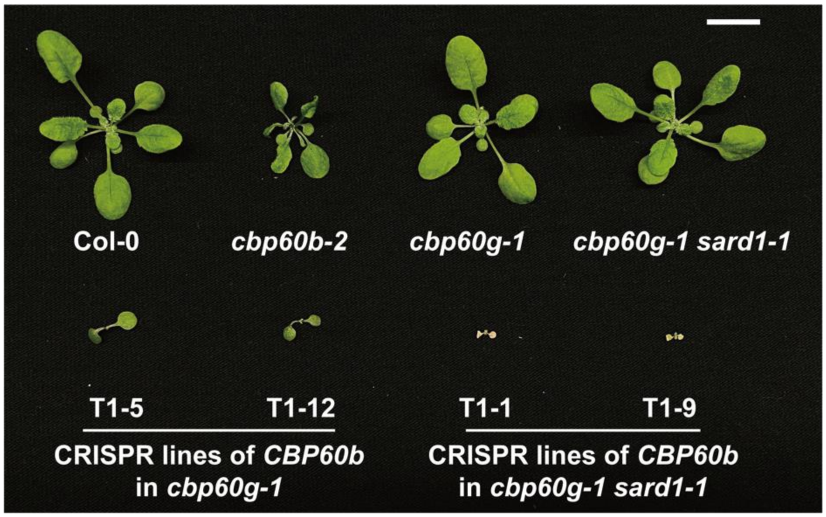


B

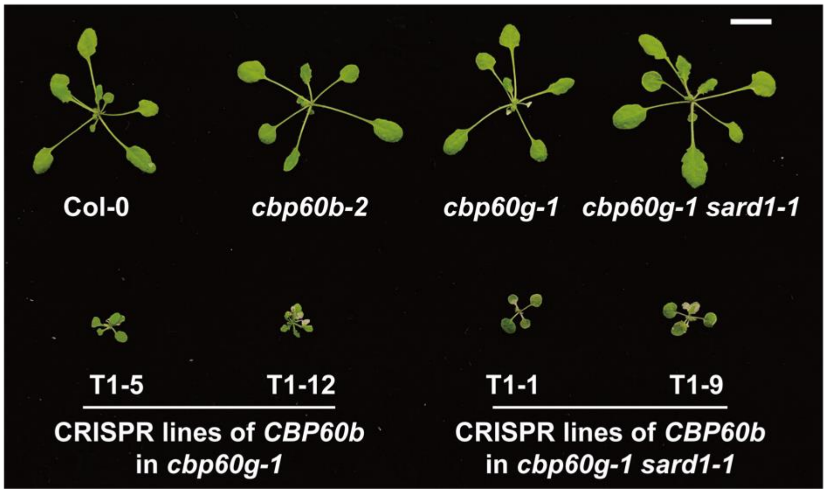


**Figure S9** Suppression of the dwarf morphologies of *cbp60b-2*, *cbp60b cbp60g-1* and *cbp60b cbp60g-1 sard1-1* by high temperature. Morphologies of three-week-old soil-grown plants of the indicated genotypes under long-day condition at 22 ºC (A) and 28 ºC (B). Scale bar is 1 cm.

### Supplementary tables

**Table S1** List of primers used in this work. Information is assorted by different applications of the primers.

Primers for quantitative PCR:

| Gene ID | Forward primer | Reverse primer |
|---|---|---|
| AT2G37620 | ACT1-F: cgatgaagctcaatccaaacga | ACT1-R: cagagtcgagcacaataccg |
| AT2G14610 | PR1-RT-F2: AGGCAACTGCAGACTCATAC | PR1-RT-R2: TTGTTACACCTCACTTTGGC |
| AT1G74710 | ICS1-F101-RT: GTCGTTCGGTTACAGGTTCC | ICS1-R102-RT: ATTAAACTCAACCTGAGGGAC |
| AT1G19250 | FMO1-F101-RT: GGAGATATTCAGTGGCATGC | FMO1-R102-RT: TTTGGTTAGGCCTATCATGG |
| AT1G73805 | SARD1-RT-NF: TCAAGGCGTTGTGGTTTGTG | SARD1-RT-NR: CGTCAACGACGGATAGTTTC |

Primers for genotyping:

| Gene ID | Forward primer | Reverse primer | Usage |
|---|---|---|---|
| AT5G57580 | AT5G57580-dele-F: gcgagtcattctcgagtttgtt | AT5G57580-dele-R: CACAACACCGACATTCCTTG | CBP60b deletion presence |
| AT5G57580 | At5g57580-homo-F: AGGAAAGAGACCGTTGCTGA | AT5G57580-dele-R: CACAACACCGACATTCCTTG | CBP60b deletion homozygosity |
| AT5G57580 | AT5G57580-dele-F: gcgagtcattctcgagtttgtt | Bda7-MT-R: TTCAGTGTTAAAGTCACCCTCAtA | cbp60g-1 presence |
| AT5G57580 | Bda7-WT-F: GCAAAGCTTCACATTGTTGTcC | AT5G57580-dele-R: CACAACACCGACATTCCTTG | cbp60g-1 homozygosity |
| AT5G26920 | CBP60g-TDNA-F: TGTTTCGGTGGACTTGTGAC | CBP60g-TDNA-R: TAAATCCCTCAACGGTCCAG | CBP60g TDNA homozygosity |
| AT1G73805 | SARD1-TDNA-F: CTTCAGTGTCGGAGTAGTCG | SARD1-TDNA-R: caagacctctctaacctaac | SARD1 TDNA homozygosity |
| AT3G48090 | EDS1-del-F: AGAACGTAAGACAGGGTTTG | EDS1-del-R: gatggagtctatattaaagagacg | EDS1 deletion presence |
| AT3G48090 | EDS1-del-F: AGAACGTAAGACAGGGTTTG | EDS1-homo-R: CATTCTCCTCTGCGAGAC | EDS1 TDNA homozygosity |

Primers for ChIP-qPCR:

| Gene ID | Forward primer | Reverse primer |
|---|---|---|
| AT1G73805 | SARD1pro0.3kb-chipF: ggaaccgtccatttgtcaac | SARD1pro0.3kb-chipR: ttcgaagaacgacaaaggaaa |
| AT1G73805 | SARD1pro-1kbF: gcacgacaagttttgagagga | SARD1pro-1kbR: agaaatgtcatgcgttaaaggaa |

Primers for cloning:

| Purpose | Primer | Sequence |
|---|---|---|
| CRISPR-deletion (CBP60b) | AT5G57580-T1-BSF | ATATATGGTCTCGATTGGAGCCAAGAGG AATTTGGAGTT |
| CRISPR-deletion (CBP60b) | AT5G57580-T2-BSR | ATTATTGGTCTCGAAACCATCCTTGCCG ATCTTGTCC |
| CRISPR-deletion (CBP60b) | AT5G57580-T1-F0 | TGGAGCCAAGAGGAATTTGGAGTTTTAG AGCTAGAAATAGC |
| CRISPR-deletion (CBP60b) | AT5G57580-T2-R0 | AACCATCCTTGCCGATCTTGTCCAATCTC TTAGTCGACTCTAC |
| pCambia1300-35S-At5g57580-3HA | AT5G57580-atgKpnI-F | ccggggtaccggtagATGATGGATAGTGGTAA |
| pCambia1300-35S-At5g57580-3HA | AT5G57580-nstopBamHi-R | CGCggatccTTCGCCATCTTCATCATCATC |
| pBasta-pAt5g57580::At5g57580-3HA | AT5G57580-KpnI-F | ccggggtaccCTGGCGTCTTGGAGTAGAGg |
| pBasta-pAt5g57580::At5g57580-3HA | AT5G57580-PstI-R | AAAActgcagTTCGCCATCTTCATCATCATC |
